# Tunable picoliter-scale dropicle formation using amphiphilic microparticles with patterned hydrophilic patches

**DOI:** 10.1101/2024.09.09.611468

**Authors:** Xinpei Song, Shreya Udani, Mengxing Ouyang, Mehmet Akif Sahin, Dino Di Carlo, Ghulam Destgeer

## Abstract

Microparticle-templated droplets or *dropicles* have recently gained interest in the fields of diagnostic immunoassays, single-cell analysis, and digital molecular biology. Amphiphilic particles have been shown to spontaneously capture aqueous droplets within their cavities upon mixing with an immiscible oil phase, where each particle templates a single droplet. Here, we fabricated an amphiphilic microparticle with four discrete hydrophilic patches embedded at the inner corners of a square-shaped hydrophobic outer ring of the particle (4C particle). 3D computational fluid dynamics simulations predicted droplet formation dynamics and differing equilibrium conditions depending on the patterning configuration. Experiments recapitulated equilibrium conditions enabling tunable dropicle configurations with reproducible volumes down to ∼200pL templated by the amphiphilic particles. The dropicle configurations depended predominantly on the size of the hydrophilic patches of the 4C particles. This validated modeling approach can inform the design of dropicles with varying volumes and numbers per particle, which can be harnessed in new amplified bioassays for greater sensitivity, dynamic range, and statistical confidence.

## 1. Introduction

Microparticle-templated droplets or *dropicles* result from either spontaneous or shear-induced partitioning of an aqueous volume within an immiscible continuous oil phase [1-8]. Dropicles have recently gained interest in the fields of diagnostic immunoassays [9-13], single-cell analysis [14-17], and digital molecular biology [18] for their ability to democratize and scale up microfluidic droplet assays that typically require complex instrumentation [19-23]. Amphiphilic microparticles, with outer hydrophobic and inner hydrophilic polymer layers, allow for spontaneous droplet formation as the aqueous phase stays within the hydrophilic cavity, reaching a stable lower energy state [1-5]. These previously reported amphiphilic microparticles [1-5], fabricated using a stop-flow lithography process [24-30], had an inner *continuous* hydrophilic layer that templated a single droplet within the particle cavity. The particle fabrication parameters were manipulated to allow for varying particle morphologies and encapsulate a broad range of droplet volumes (0.25nL to 30nL) [3,5]. These dropicles with volumes *O*(nL) were used to detect analytes with a detection limit of 10pg/ml in buffer [2] and 50pg/ml in serum [4]. For enhanced bioassay sensitivity and accuracy, it is essential to reduce the volume of segmented droplets and increase the number of individual compartments per assay for better statistical confidence. One way to achieve this is to fabricate smaller amphiphilic particles with even tinier cavities to hold droplets with volumes *O*(pL). However, such an approach is plagued with challenges on the fabrication end, such as difficulty stopping the flow in smaller microfluidic channels, as depicted in our earlier work [3]. Therefore, an out-of-the-box approach is required to fabricate new particle designs that can hold smaller droplet volumes *O*(pL) without significantly reducing the particle size.

The dropicle formation depends on the amphiphilic particle’s shape and polymer distribution, which is defined by a 3D-printed microfluidic device that could sculpt multiple co-flowing streams in a target cross-sectional shape [2-4]. However, it is costly and time-consuming to experimentally vary the design of the microfluidic devices for modulating the characteristics of the fabricated particle. Therefore, the first step towards the desired dropicle formation by optimizing the particle parameters is to meticulously design the microfluidic devices using numerical models. Numerical simulations offered an indispensable way to predict the particle’s characteristics before the 3D printing of the microfluidic device and fabricating the particles. We have previously used single-phase flow simulations to predict the shape of the sculpted flow profile inside microfluidic channels [2]. However, these numerical models do not account for the variable viscosities of different co-flowing streams. Therefore, a two-phase flow model is required to simulate variable viscosity co-flowing streams and obtain an accurate sculpted flow cross-section before particle fabrication.

Moreover, numerical modeling of dropicle formation before experimental testing of new particle designs could save precious lab resources and time. To this end, mathematical models minimizing the interfacial energy are introduced to simulate uniform droplet volumes templated by single-material crescent or multi-material cylindrical particles [31,32]. Recently, we have established and analyzed a 2D computational fluid dynamic (CFD) model to simulate the interface of immiscible water and oil phases for dropicle formation within concentric amphiphilic particles [33]. The numerical findings matched well with the experimental results [3]. A 2D CFD model can be suitable for symmetric particle geometries, e.g. O-shaped particles, with a reasonable computational cost. However, it cannot encompass complex particle shapes with anisotropic material distribution and asymmetric dropicle formation. Therefore, a 3D CFD model is required to understand the formation of dropicles in various shapes dictated by the shape of the amphiphilic particle.

In this work, we demonstrate amphiphilic particle designs with four discrete and carefully patterned hydrophilic regions embedded within the four corners of a square-shaped hydrophobic backbone (4C particles). We study, computationally and experimentally, the ability of 4C particles to form picoliter-scale localized droplets. We designed and 3D printed a multi-channel microfluidic device to sculpt a multi-layered laminar flow of polymer precursors and fabricate the 4C particles using a ‘stop flow lithography’ process (**Figure 1A** and 1B). We performed two-phase flow simulations to predict the shape of the particles by the sculpted flow profile, defined by variable viscosity polymer precursor streams, within the outlet channel of the device. The fabricated 4C particles with tunable physio-chemical characteristics were later used to template one to four droplets per particle of varying volumes *O*(pL) and morphologies, such that a single square droplet (S^1^), a triangular droplet accompanied by one small droplet (T^+1d^), a rectangular droplet accompanied by two small droplets (R^+2d^) or four individual droplets at the corners (C^4d^) were captured by the 4C particles (Figure 1C and 1D). We also developed a 3D numerical model to describe the formation of variable dropicle configurations and study how the dropicle shape can evolve over time due to particle characteristics, swelling, and trapped aqueous volume.

**Figure 1.**
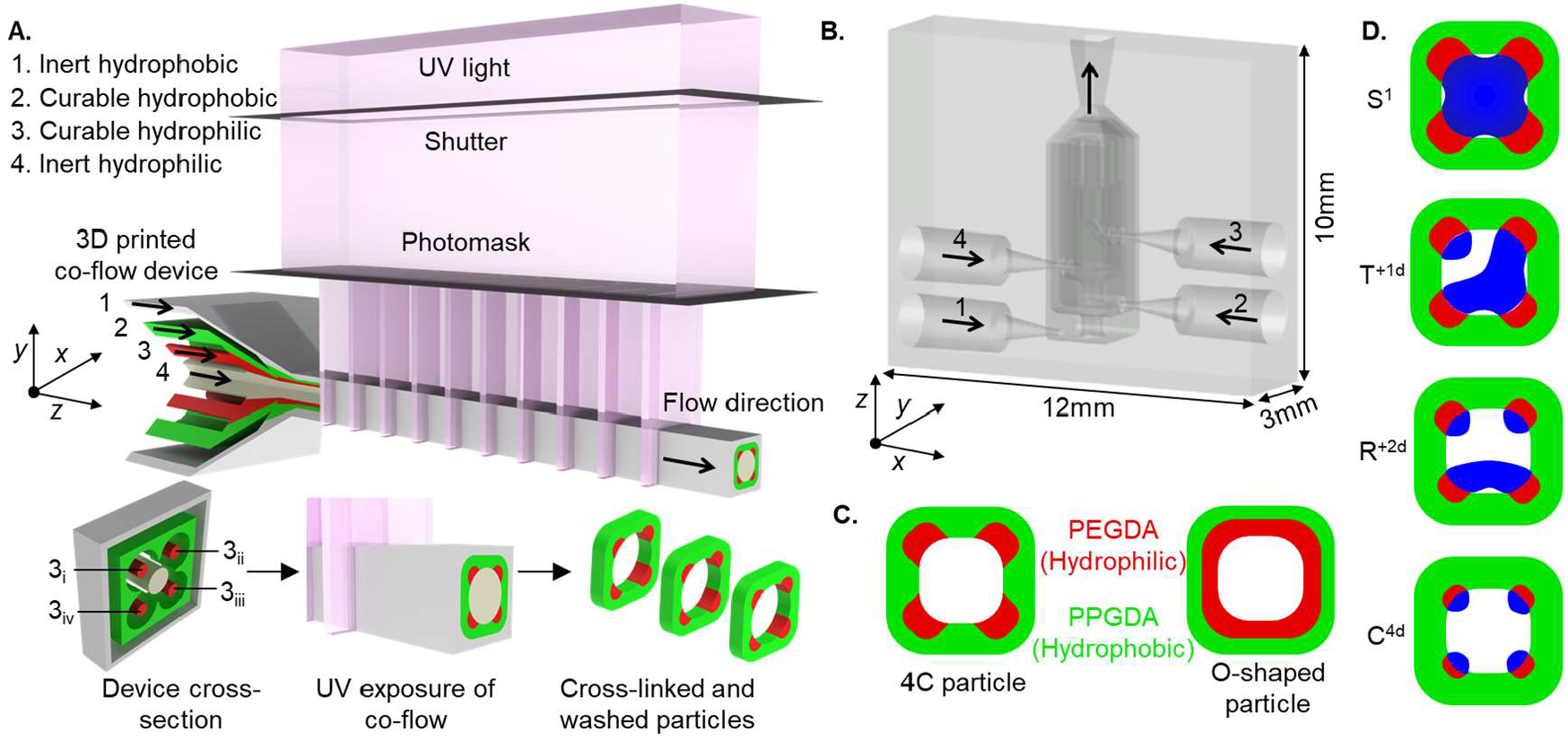
(A) Schematic of the 3D printed microfluidic device for 4C particle fabrication using the stop-flow lithography process. (B) A to-the-scale model of the 3D microfluidic device. (C) Comparison between the 4C and O-shaped particles. (D) Four different dropicle configurations are driven by the tunable design of 4C particles.

## 2. Results and Discussions

### 2.1 4C particle fabrication and characterization

We performed a 3D numerical simulation of fluid flow through the microfluidic device (quarter domain) to predict the cross-sectional profile of the sculpted flow (**Figure 2A**). The green and red color streamlines indicate the curable hydrophobic and hydrophilic precursor streams, respectively, that will define the particle shape. The flow rate ratios of the four streams *Q*_1,2,4_:*Q*_3_ were varied as 1:1, 2:1, 4:1, 9:1, 12:1, and 16:1, to obtain F_1_, F_2_, F_4_, F_9_, F_12_, and F_16_ particles, respectively. The flow rate ratio was increased while maintaining a constant total flow rate (*Q*_T_ = *Q*_1_+*Q*_2_+*Q*_3_+*Q*_4_) to gradually decrease the hydrophilic patch size of the 4C particles from F_1_ to F_16_ (Figure 2B). The particle shape predicted by the CFD simulation matched with the cured particles suspended in ethanol (EtOH); however, minor discrepancies between the experimental and numerical results could be attributed to the fact that the cured particles deformed slightly after they were transferred from a polymer precursor solution to pure EtOH solution (see also **Figure S1**).

**Figure 2.**
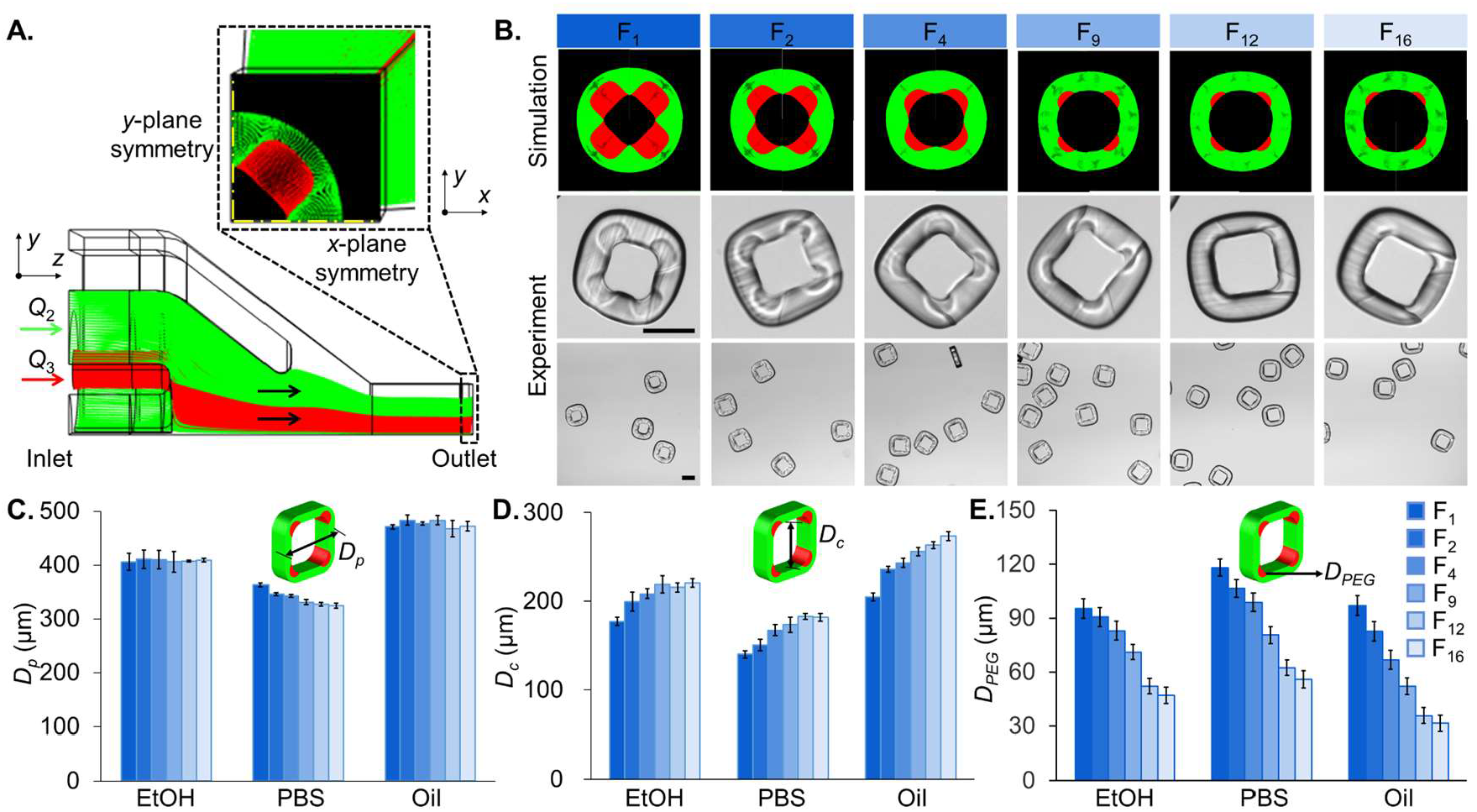
Particle fabrication and characterization. (A) Quarter 3D model of the microfluidic device to generate the particle. (B) Simulated and experimental particles with different flow rate ratios. The top row shows the simulated particles in the form of velocity streamlines at the outlet of the microchannels. The following two rows are the experimentally observed particles, with a zoomed view of a single particle (middle) and multiple particles (bottom) in EtOH. Scale bar: 200µm. (C-E) Distributions of *D*_*p*_, *D*_*c*_, and *D*_*PEG*_ in EtOH, PBS, and oil with different flow rate ratios. The inserts are the schematic of the particle.

The medium around particles is sequentially changed from EtOH to phosphate-buffered saline (PBS) to oil for dropicle formation (section 2.2). The particle (*D*_*p*_), cavity (*D*_*c*_), and hydrophilic patch (*D*_*PEG*_) diameters measured in EtOH, PBS, and oil showed reasonable uniformity within each particle type (Figure 2C-E). The 4C particles consistently shrank from EtOH to PBS and expanded from PBS to oil. In EtOH, all the particle types (F_1_ to F_16_) showed a narrow size distribution with *D*_*p*_ of 409 ± 3µm, even though we used very different flow rate ratios during the fabrication process, which highlights the consistency of the backbone structure of the particles (Figure 2C). However, upon medium exchange from EtOH to PBS, each particle type contracted differently following the hydrophobic to hydrophilic polymer ratio within a particle. The F_1_ particle, with the lowest proportion of hydrophobic polymer, shrank the least from *D*_*p*_ ≈ 406µm to ≈ 363µm (∼11%), whereas F_16_ particles, with the highest proportion of hydrophobic polymer, shrank the most from *D*_*p*_ ≈ 409µm to ≈ 324µm (∼21%). As the medium was exchanged from PBS to oil, all the particle types (F_1_ to F_16_) expanded beyond their original sizes in EtOH, with an average *D*_*p*_ ≈ 475 ± 7µm. For example, the F_1_ and F_16_ particles expanded back by ∼30% and ∼46% from PBS to oil, respectively. A higher expansion of F_16_ particles can be attributed to the soaking of a relatively thicker hydrophobic layer by oil. A variable expansion of F_1_-F_16_ particles in oil contributes to different dropicle configurations in subsequent experiments (sections 2.2-2.4).

For particles F_1_ to F_16_, the cavity diameter in EtOH increased by ∼25% from *D*_*c*_ ≈ 177µm to ≈ 221µm (Figure 2D). For any given particle type, the *D*_*c*_ decreased from EtOH to PBS and increased back in oil, following a trend similar to *D*_*p*_. For example, the F_1_ particle cavity shrunk by ∼21% from EtOH to PBS and increased by ∼46% from PBS to oil. With a gradual increase in the flow rate ratio, *Q*_1,2,4_:*Q*_3_, from 1:1 to 16:1, the hydrophilic proportion within a particle receded, which left space for the innermost inert stream *Q*_*4*_ to expand the cavity diameter. The hydrophilic patch diameter in EtOH also decreased significantly from *D*_*PEG*_ ≈ 95µm to ≈ 47µm (i.e., ∼51% reduction) as the flow rate ratio was increased from F_1_ to F_16_ (Figure 2E). Contrary to the particle and cavity diameters (*D*_*p*_ and *D*_*c*_), the hydrophilic patch (*D*_*PEG*_) expanded from EtOH to PBS and shrank in oil. For example, in the F_1_ particles with *D*_*PEG*_ ≈ 95µm in EtOH, the PBS was absorbed by the hydrophilic patch to swell it to *D*_*PEG*_ ≈ 118µm (i.e., ∼24% increase). The oil phase pushed the PBS to a minimal surface energy configuration, resulting in the contraction of the hydrophilic patch with *D*_*PEG*_ ≈ 97µm (i.e., ∼18% reduction). A variable hydrophilic patch size for particles F_1_ to F_16_ was the most critical feature of the particles for variable dropicle formations.

### 2.2 Numerical modeling of dropicle formation

An aqueous droplet is captured within the cavity of the 4C particle as the media is exchanged from EtOH to PBS to oil (**Figure 3A**). We established a two-phase flow 3D CFD model, based on our earlier reported 2D model [33], to systematically investigate the mechanism driving different dropicle configurations within the particle cavities (Figure 3B-D). The dropicle configuration was influenced by the hydrophilic patch radii *R*_1-4_ and the ratio of the droplet initial height (*H*_*d*_) with respect to the particle cavity height (*H*_*c*_). We modeled dropicle formation using orthotropic, symmetric, and asymmetric particle geometries for a range of *H*_*d*_/*H*_*c*_ values and a fixed simulation time (Figure 3B). The *H*_*d*_/*H*_*c*_ ratio corresponds to the ratio of the droplet to the particle cavity volumes (*V*_*d*_/*V*_*c*_), which varied from 0.75 to 0.25, assuming the cavity was never filled to its maximum capacity due to cavity expansion from PBS to oil (see Figure 2D). Notably, a ∼50% increase in *D*_*c*_ from PBS to oil will expand the cavity volume by >2x. Therefore, it is reasonable to assume that the final cavity volume *V*_*c*_ will always be larger than the volume of the droplet *V*_*d*_ it encompasses. Hence, we performed the numerical simulations for *H*_*d*_/*H*_*c*_ or *V*_*d*_/*V*_*c*_ ranging from 0.75 to 0.25.

**Figure 3.**
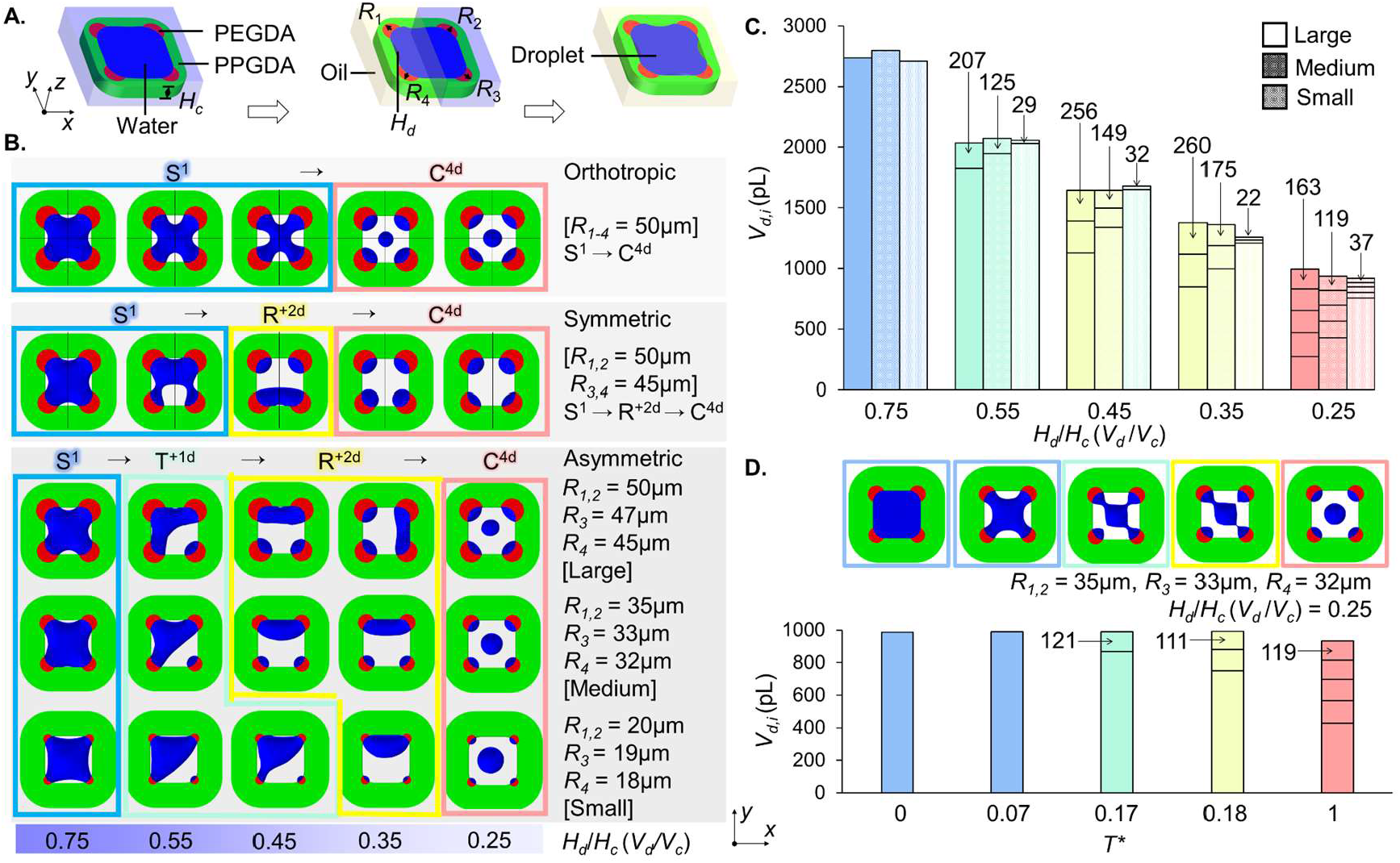
(A) Schematic of the dropicle formation within a 4C amphiphilic particle upon medium exchange from EtOH to PBS to Oil. (B) Three numerical models, orthotropic, symmetric, and asymmetric, are solved to systematically obtain different dropicle configurations for variable *H*_*d*_/*H*_*c*_ ratio. In the asymmetric model, the hydrophilic patch radii (*R*_1-4_) are varied from large to medium to small combinations to realize the dropicle transition from S^1^ → T^+1d^ → R^+2d^ → C^4d^ for variable *H*_*d*_/*H*_*c*_ ratio. (C) A quantitative analysis of the individual droplet volumes (*V*_*d,i*_) for different dropicle configurations and *H*_*d*_/*H*_*c*_ ratio. The stacked bars represent the volume of each droplet in configurations with multiple droplets per particle. (D) A time-dependent simulation highlights the dropicle transition over time as the total aqueous volume is split up into smaller volumes *V*_*d,i*_.

The orthotropic particle, with four identical patches at the corners, formed an S^1^ droplet for *H*_*d*_/*H*_*c*_ = 0.75, which transitioned directly to C^4d^ droplets adhering to the four hydrophilic patches of the particle for *H*_*d*_/*H*_*c*_ ≤ 0.35 (Figure 3B). An orthotropic model can not produce the T^+1d^ or R^+2d^ configurations. The symmetric particle, with two pairs of identical hydrophilic patches, resulted in a similar S^1^ droplet for *H*_*d*_/*H*_*c*_ = 0.75, which transitioned to R^+2d^ and then C^4d^ droplets as the *H*_*d*_/*H*_*c*_ decreased to 0.45 and 0.35, respectively. The symmetric model is unable to produce the T^+1d^ configuration. However, an asymmetric particle, with three distinct hydrophilic patch radii, encompassed all the dropicle configurations transitioning as follows: S^1^ → T^+1d^ → R^+2d^ → C^4d^, as *H*_*d*_/*H*_*c*_ decreased from 0.75 to 0.25. Notably, a difference of ≲10% between the hydrophilic patch radii *R*_1-4_ was enough to induce asymmetric dropicle shapes, i.e., T^+1d^ and R^+2d^. It was essential to model the dropicle formation with an asymmetric particle design to capture all four types of dropicle configurations.

We simulated dropicle formation using particles with large, medium, and small radii combinations [*R*_1,2_, *R*_3_, *R*_4_] of [50, 47, 45 µm], [35, 33, 32 µm], and [20, 19, 18 µm], respectively, for variable *H*_*d*_/*H*_*c*_ or *V*_*d*_/*V*_*c*_ values (Figure 3B and **Movie S1**). We selected different radii *R*_1-4_ values within a single particle to model the slight variability between the hydrophilic patches due to uneven or decaying UV exposure within the microchannel during the particle fabrication. For all asymmetric particle models, we obtained a gradual transition of dropicle configurations from S^1^ → T^+1d^ → R^+2d^ → C^4d^ as the *H*_*d*_/*H*_*c*_ decreased from 0.75 to 0.25. For the small radii combination, the dropicle transitioned from T^+1d^ → R^+2d^ as the *H*_*d*_/*H*_*c*_ decreased from 0.45 to 0.35, whereas the R^+2d^ configuration could only be observed for a very narrow range of *H*_*d*_/*H*_*c*_ values. The large hydrophilic patches exert a relatively greater pull on the aqueous droplet towards the corners due to the wettability of the patches for a constant PBS-oil interfacial tension. Therefore, the dropicle transitioned early from T^+1d^ → R^+2d^ as the PBS volume fraction (*V*_*d*_/*V*_*c*_) decreased below 0.55 for large and medium patches. For the small patches, the wetting force pulling the droplet toward the corners was not strong enough; therefore, the T^+1d^ configuration was retained for 0.55 ≥ *V*_*d*_/*V*_*c*_ ≥ 0.45. However, a further reduction in droplet volume resulted in a transition to R^+2d^ dropicles for *V*_*d*_/*V*_*c*_ = 0.35 and finally to C^4d^ dropicles for *V*_*d*_/*V*_*c*_ = 0.25. If we associate the large patch radii with higher droplet volume (i.e., *V*_*d*_/*V*_*c*_ = 0.75) retained within the cavity, one can easily deduce that the most stable dropicle configuration will be S^1^. On the contrary, the small patch radii with lower droplet volume (i.e., *V*_*d*_/*V*_*c*_ = 0.25) retained within the cavity will consistently result in the C^4d^ dropicles. The other configurations, i.e., T^+1d^ and R^+2d^, will fall in between the large and small patch radii.

A quantitative analysis of dropicle formation revealed that the 4C particles with *D*_*c*_ of 200μm can form droplets with a wide range of volumes ranging from >2.5nL to <100pL (Figure 3C). The 4C particles, irrespective of hydrophilic patch size, formed single S^1^ dropicles with volumes of ∼2.75nL for *V*_*d*_/*V*_*c*_ of 0.75. For *V*_*d*_/*V*_*c*_ = 0.55, the dropicles transitioned from S^1^ to T^+1d^ configuration. For the large patch particle, we obtained the T^+1d^ configuration with two droplets of ∼207pL and ∼1.80nL volume, respectively. The medium and small patch particles were able to form even smaller droplets of ∼125pL and ∼29pL, respectively, at one corner of the particle cavity with T^+1d^ configuration, which can be attributed to the reduced patch sizes. For *V*_*d*_/*V*_*c*_ = 0.45, the 4C particles with large and medium patches formed dropicles with R^+2d^ configuration with the volume of the smallest droplets as ∼256pL and ∼149pL, respectively. The particles with small patches could still retain two T^+1d^ dropicles with volumes of ∼1.65nL and ∼32pL. For *V*_*d*_/*V*_*c*_ = 0.35, all the particles resulted in an R^+2d^ configuration. Similarly, for *V*_*d*_/*V*_*c*_ = 0.25, all the particles resulted in the C^4d^ configuration with four droplets formed at the four corners of the cavity and a satellite droplet at the center. The average volumes of the four C^4d^ droplets were ∼181pL, ∼127pL, and ∼41pL for the large, medium, and small patch particles, respectively. It indicated that we can obtain smaller droplets within the 4C amphiphilic particle by adjusting the patch size without changing the particle size. A small variation in total aqueous volume between simulations can arise from the numerical error. Moreover, a droplet diameter below a critical value can shrink due to rapid evaporation of the aqueous phase in the numerical models, leading to the variation in overall aqueous volume between the simulations [34, 35].

Time-dependent numerical models enabled us to analyze the dropicle shape transition over a normalized time scale *T*\* (Figure 3D). For medium-sized hydrophilic patches and *V*_*d*_/*V*_*c*_ of 0.25, the droplet shape transitioned from S^1^ → T^+1d^ → R^+2d^ → C^4d^ at *T\** = 0.07, 0.17, 0.18, and 1.00, respectively. The total aqueous volume within the cavity remained constant at ∼990pL, which gradually split up from a single droplet to four isolated droplets at the corners of the particle with volumes as low as ∼119pL.

### 2.3 Experimental analysis of dropicle formation

We experimentally investigated the dropicle formation using different 4C particles (F_1_-F_16_) as the hydrophilic patches of variable sizes retained matching PBS volumes within the particle cavities, sealed by the immiscible oil phase (**Figure 4A**). The 4C particles, F_1_, F_9_, F_12_, and F_16_, resulted in a dominant dropicle configuration of S^1^, T^+1d^, R^+2d^, and R^+2d^+C^4d^, respectively, as depicted in the bright field and fluorescent images of the dropicles. The experimental results agreed well with the corresponding numerical models built using different *V*_*d*_/*V*_*c*_ ratios and hydrophilic patch radii combinations (*R*_1,2_, *R*_3_, *R*_4_), which represented the F_1_, F_9_, F_12_, and F_16_ particles, respectively (Figure 4B). The average patch diameters of 95, 71, 52, and 47µm measured experimentally for the F_1_, F_9_, F_12_, and F_16_ particles were similar to the patch diameters in the numerical simulations 96, 69, 48, and 39µm. The experimental S^1^ dropicles within F_1_ particles corresponded to the large hydrophilic patches that captured the highest aqueous volume during the dropicle formation, which was numerically modeled with *V*_*d*_/*V*_*c*_ = 0.65 to result in the same S^1^ dropicle configuration. As the hydrophilic patches got smaller from F_1_ to F_16_ particles, the capability of holding aqueous volume within the cavities decreased. Therefore, the numerical models for the F_9_, F_12_, and F_16_ particles were established using *V*_*d*_/*V*_*c*_ of 0.48, 0.35, and 0.28, which resulted in the T^+1d^, R^+2d^, and C^4d^ configurations, respectively. In the experiments, the dominant dropicle configurations for the F_9_, F_12_, and F_16_ particles were T^+1d^, R^+2d^, and R^+2d^+C^4d^, which aligned well with the numerical results.

**Figure 4.**
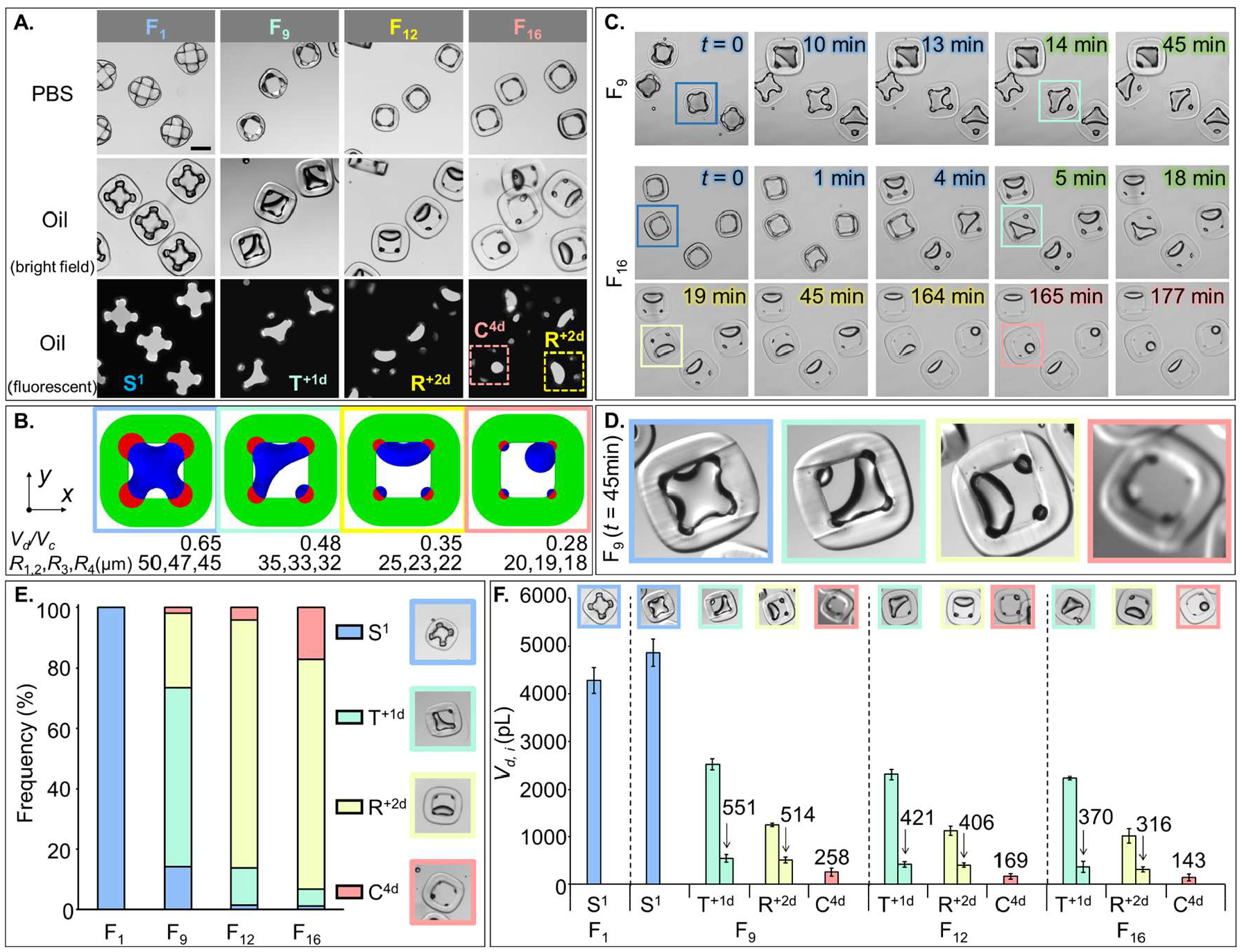
(A) Experimental observation of the dropicle formation within F_1_, F_9_, F_12_, and F_16_ particles. Upon addition of the oil phase, aqueous PBS dropicles (visible in the brightfield and fluorescent images) were captured within the particle cavities. (B) Numerical models of the dropicle formation within particles with variable hydrophilic patch radii match well with the experimental results in (A). (C) Transition of dropicle configurations over time within F_9_ (S^1^ → T^+1d^) and F_16_ (S^1^ → T^+1d^ → R^+2d^ → C^4d^) particles. (D) A close-up view of a single F_9_ particle with four dropicle configurations reached at *t* = 45min. (E) Frequency distributions of S^1^, T^+1d^, R^+2d^, and C^4d^ dropicles within F_1_, F_9_, F_12_, and F_16_ particles. (F) The average volume of individual droplets *V*_*d,i*_ within the S^1^, T^+1d^, R^+2d^, and C^4d^ dropicles captured by the F_1_, F_9_, F_12_, and F_16_ particles.

A dropicle formed within a given 4C particle can transition from one configuration to the next over time (Figure 4C, **Movie S2** and **Movie S3**). For example, S^1^ dropicles formed inside the F_9_ particles at *t* = 0min gradually transitioned to a stable T^+1d^ configuration at *t* = 45min. We could observe a complete transition of dropicle configurations from S^1^ → T^+1d^ → R^+2d^ → C^4d^ within F_16_ particles observed for ∼3h. The transition from S^1^ → T^+1d^ → R^+2d^ occurred within the first 19 min after dropicle formation. The R^+2d^ remained the most dominant and stable configuration from *t* = 19 min to 45 min even though the aqueous droplets kept shrinking due to gradual evaporation over time. At *t* ≥ 164 min, we see the emergence of the C^4d^ configuration in addition to the R^+2d^. A single particle type can exhibit more than one dropicle configuration based on particle-to-particle variability during fabrication and a slightly different aqueous volume captured within the particles during the dropicle formation. For example, at a given time *t* = 45 min after dropicle formation, the F_9_ particles depicted all four of the dropicle configurations, i.e., S^1^, T^+1d^, R^+2d^, and C^4d^, at variable frequencies; however, the T^+1d^ configuration remained the dominant one (Figure 4D and 4E).

We quantitatively characterized different dropicle configurations, i.e., S^1^, T^+1d^, R^+2d^, and C^4d^, formed within the F_1_, F_9_, F_12_, and F_16_ particles by plotting their frequency distributions (Figure 4E). The F_1_, F_9_, F_12_, and F_16_ particles resulted in predominantly S^1^ (100%), T^+1d^ (59%), R^+2d^ (82%), and R^+2d^ (76%) dropicles, respectively. The proportion of S^1^ dropicles dropped to 14% for F_9_ particles and <2% for F_12_ and F_16_ particles. Similarly, the proportion of T^+1d^ decreased from 59% (F_9_) to 13% (F_12_) and 6% (F_16_). However, the R^+2d^ dropicle frequency increased from 25% (F_9_) to 82% (F_12_) and 76% (F_16_). The highest proportion of C^4d^ dropicles (17%) was obtained within the F_16_ particles, which was significantly higher than the C^4d^ proportion in F_9_ (2%) and F_12_ (4%) particles. The data for F_2_ and F_4_ dropicles were included in **Figure S3**.

Moreover, we plotted the average volume of individual droplets *V*_*d,i*_ captured inside the 4C particles (Figure 4F). The average S^1^ dropicle volume calculated for F_1_ particles was ∼4.3nL, which slightly increased to ∼4.9nL for F_9_ particles due to an increase in their cavity diameter *D*_*c*_. The F_9_ particles also formed T^+1d^, R^+2d^, and C^4d^ dropicles, where the smallest average dropicle volume was recorded for C^4d^ configuration at ∼258pL. The number of S^1^ dropicles within F_12_ and F_16_ particles was negligible, therefore, the S^1^ volumes were not plotted for these particles. The volumes of individual droplets within T^+1d^, R^+2d^, and C^4d^ configurations decreased further for the F_12_ and F_16_ particles, confirming the theory that smaller hydrophilic patches retain lower droplet volume. We measured the smallest average volume of ∼143pL for the C^4d^ dropicle captured within F_16_ particles.

### 2.4 Comparison between amphiphilic particles with continuous and discrete hydrophilic layers

To analyze the influences of the continuous and discrete hydrophilic layers on the dropicle formation within an amphiphilic particle, we compared the particle dimensions and dropicle configurations between the O-shaped and 4C amphiphilic particles with *D*_*p*_ = ∼400µm (**Figure 5**). The experimental data for the O-shaped particle was adapted from our previous work [3]. We calculated the contraction (-) and expansion (+) rates (Δ) of the measured particle diameter *D*_*p*_, cavity diameter *D*_*c*_, hydrophilic layer thickness *T*_*PEG*_ (*O*-shaped particle), and hydrophilic patch diameter *D*_*PEG*_ (4C particle), for medium exchange from EtOH → PBS and from PBS → oil, respectively (Figure 5A). All the dimensions of the 4C particle changed less than that of the O-shaped particle during the media exchange. For EtOH → PBS exchange, the O-shaped particles shrank relatively more with -16% Δ*D*_*p*_ and -32% Δ*D*_*c*_, compared to the 4C particles, having an inner discontinuous hydrophilic with -11% Δ*D*_*p*_ and -21% Δ*D*_*c*_. For PBS → oil exchange, the Δ*D*_*p*_ and Δ*D*_*c*_ values for the 4C particle, i.e. +30% and +46%, were smaller than that of the O-shaped particle, i.e. +37% and +58%. For PBS → oil exchange, the Δ*T*_*PEG*_ of - 30% for the O-shaped particle was clearly higher than the Δ*D*_*PEG*_ of -18% for the 4C particle.

**Figure 5.**
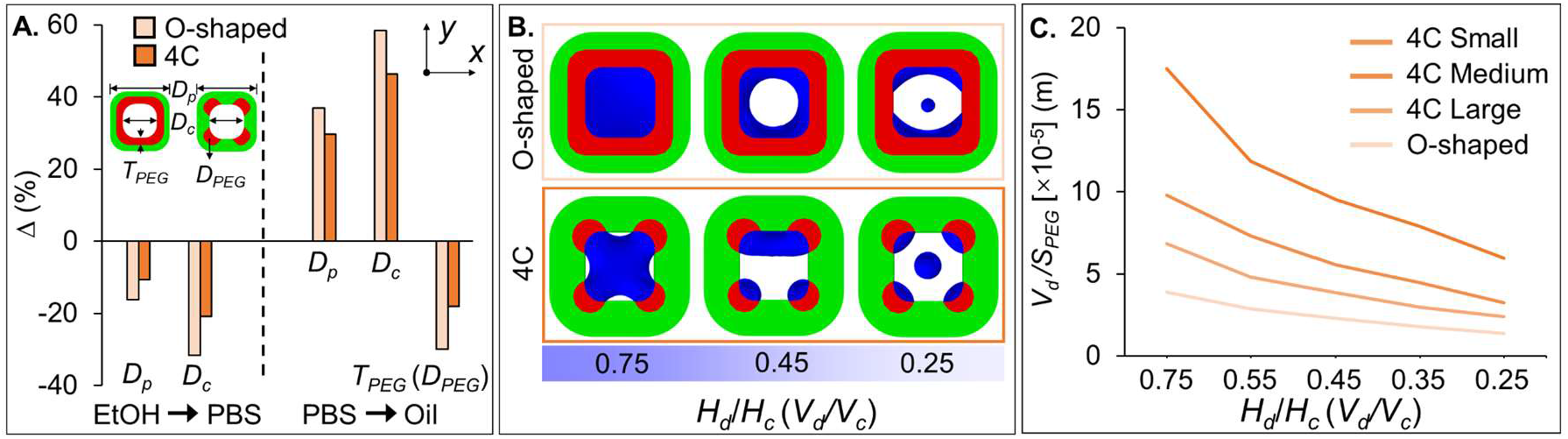
Comparison between the O-shaped and 4C particles. (A) The contraction and expansion rates of the particle diameter *D*_*p*_, cavity diameter *D*_*c*_, thickness *T*_*PEG*_ (O-shaped particle), and diameter *D*_*PEG*_ (4C particle) of the hydrophilic layer in EtOH, PBS, and oil. (B) Simulated dropicle formation within the O-shaped and 4C particles (large hydrophilic patches) with different *H*_*d*_/*H*_*c*_ values. Here, *D*_*p*_ = 400µm, *D*_*c*_ = 200µm, *T*_*PEG*_ = 50µm, and *R*_1,2_, *R*_3_, *R*_4_ = 50µm, 47µm, 45µm. (C) Total droplet volume *V*_*d*_ to hydrophilic surface area *S*_*PEG*_ ratio for the O-shaped and 4C dropicles with different hydrophilic patch sizes.

We further simulated the aqueous volume distributions within the O-shaped and 4C particles with varying *V*_*d*_/*V*_*c*_ values (Figure 5B). A droplet filled the cavity of the O-shaped particle entirely when *V*_*d*_/*V*_*c*_ = 0.75. However, the cylindrical droplet transitioned to a single annular ring-shaped droplet gradually at *V*_*d*_/*V*_*c*_ = 0.45. For *V*_*d*_/*V*_*c*_ = 0.25, two separated droplets were formed adhering to the inner hydrophilic layer of the particle accompanied by a satellite droplet at the cavity center. Notably, the hydrophilic surface wettability dominated the force balance with the water-oil interfacial tension to result in a thinner ring-shaped droplet or two segregated droplets clinging to the inner layer of the particle. For a stable circular droplet obtained experimentally [3], the O-shaped particles should capture a sufficient volume of water inside its cavity, which corresponds to *V*_*d*_/*V*_*c*_ = 0.75. Comparatively, the 4C dropicles resulted in very different configurations and multiple droplets driven by the discrete hydrophilic patches as described above. Lastly, we have plotted the total dropicle volume (*V*_*d*_) to hydrophilic wetting surface area (*S*_*PEG*_) ratio (*V*_*d*_/*S*_*PEG*_) for the 4C and O-shaped particles (Figure 5C). The *V*_*d*_/*S*_*PEG*_ ratio increased by ∼1.8x, ∼2.5x, and ∼4.4x for the 4C particles with large, medium, and small hydrophilic patches, respectively, when compared with the O-shaped particles. Such ability to vary the *V*_*d*_/*S*_*PEG*_ ratio can enable new bioassay development with tunable sensitivities and dynamic ranges.

## 3. Methods

### 3.1 Numerical modeling

#### 3.1.1 Simulation for particle shape prediction

To predict the particle’s shape and dimensions under different flow rate ratios, we adopted the “Two-Phase Flow, Level Set” module in COMOSL Multiphysics 6.1 to run laminar flow streams of variable velocities through an outlet microchannel with a dimension of 0.5mm. To reduce the computing time, we established a quarter model of the microchannel (Figure 2A). The interfacial tension between the two streams was set as zero to emulate minimal free surface energy between the partially miscible streams. The viscosities of the inner hydrophilic and outer hydrophobic streams were set to be 0.007Pa·s and 0.042Pa·s, respectively. The density of the two flow streams was constant, 987kg/m^3^. The contact angle of the interface with the wetted walls of the microchannel was 90°. The flow rate ratio between the four streams *Q*_1_:*Q*_2_:*Q*_3_:*Q*_4_ was altered to obtain particle shapes with variable thicknesses of the discrete hydrophilic patches and the continuous hydrophobic layer of the particle. The total flow rate for the four inlets was 2mL/min. We also solved the “Laminar Flow” module to compare the results of the “Two-Phase Flow, Level Set” module with a single-phase flow model (SI).

#### 3.1.2 Simulation for dropicle formation

We used the “Two-Phase Flow, Phase Field” module to simulate the various dropicle configurations formed inside the particle cavity. For optimal computing time, we established a vertically symmetric half model. An amphiphilic particle was modeled containing a continuous outer hydrophobic layer and four hydrophilic patches at the corners of the particle. The cavity of the particle was uniformly filled with an aqueous solution at the onset of the simulation, whereas the oil phase surrounded the particle. The interfacial tension between the water and oil phases was 0.03N/m. The contact angles that the interface would make with the hydrophilic and hydrophobic layers of the particle were fixed at 45° and 150°, respectively. The density and dynamic viscosity of the aqueous phase were 1000kg/m^3^ and 0.001Pa·s, and that of the oil phase were 1050kg/m^3^ and 0.06Pa·s, respectively. We employed an adaptive mesh, where a finer mesh region followed the water-oil interface as the simulation moved forward in time (**Figure S2**). The water-oil interface thickness was set as 10x smaller than the default thickness value to trace the interface accurately. The simulation output time was *T* = 10ms (Figures 3B-C and 4B). To evaluate the volume of individual droplets within a given 4C dropicle configuration, we partitioned the cavity domain into 16 smaller portions for the appropriate estimation of each droplet volume (Figure 3D-E). For the O-shaped dropicle formation, the simulation conditions were identical to that of the 4C dropicle (Figure 5B).

### 3.2 Experimental

#### 3.2.1 Particle fabrication using stop-flow lithography

We introduce four polymer precursor streams, i.e., inert hydrophobic, curable hydrophobic, curable hydrophilic, and inert hydrophilic, with different flow rates *Q*_1_-*Q*_4_ from the inlets 1-4 of the 3D printed microfluidic device (Figure 1A). The inlets open into four stacked microchannels separated by thin walls to prevent the premature mixing of streams (Figure 1B). Notably, inlet 3 was further divided into four separated yet identical subchannels to bring four uniform streams (3_*i*_-3_*iv*_) of the curable hydrophilic precursor to the device. The four main streams 1-4 flowed along their respective microchannels and sequentially merged at a tapered region of the device close to the outlet to enable a multi-layered structured co-flow of hydrophilic and hydrophobic precursors inside a square glass capillary. The curable precursor streams, mixed with a photo-initiator (PI), are polymerized under a photomask to form amphiphilic microparticles with discontinuous inner hydrophilic patches and a continuous outer hydrophobic layer. The rectangular slits in the photomask define the particle height (*H*_*p*_), whereas the flowrates *Q*_*1*_*-Q*_*4*_ influence the particle (*D*_*p*_), cavity (*D*_*c*_), and hydrophilic patch (*D*_*PEG*_) diameters. Compared to a concentric O-shaped amphiphilic particle with a continuous inner hydrophilic layer [2,3,5], the 4C particle has four independent hydrophilic patches at the corners (Figure 1C). The particles were collected in the EtOH solution for the subsequent dropicle formation.

#### 3.2.2 Dropicle formation

The workflow to form the particle-templated droplets within the 4C amphiphilic particle was based on simple pipetting and washing steps (Figure 3A). Firstly, the particles suspended in EtOH were transferred to a well plate, followed by a medium exchange to a PBS solution. The hydrophilic patches swelled after absorbing the aqueous PBS solution, whereas the outer hydrophobic layer contracted. Secondly, the excess PBS was removed, and the oil (poly(dimethylsiloxane-co-diphenylsiloxane)) was added to the well plate to push away the aqueous phase outside the particles due to the immiscibility between the water and oil phases. The droplets inside each 4C amphiphilic particle were encapsulated by the continuous oil phase.

#### 3.2.3 Dropicle volume calculation

We multiplied the droplet area by *H*_*c*_ and a factor (*H*_*d*_/*H*_*c*_) to derive the droplet volume for the S^1^, T^+1d^, R^+2d^, and C^4d^ dropicles (Figure 4F). The droplet area of each shape was measured using Fiji software. The factors (*H*_*d*_/*H*_*c*_) were adopted from numerical simulation (Figure 4B) as 0.65, 0.48, 0.35, and 0.28, for S^1^, T^+1d^, R^+2d^, and C^4d^ dropicles, respectively.

## 4. Conclusions

We have proposed 4C amphiphilic particles composed of an outer continuous hydrophobic layer embedded with four inner discrete hydrophilic patches at the corners, where the patch size could be readily tuned by modulating the flow rate ratio during particle fabrication. The 4C particles fabricated with different patch sizes were able to retain dropicles in four different configurations. We have conducted systematic numerical simulations for particle shape prediction and dropicle formation within the particle cavity. We have experimentally measured and characterized the particle dimensions in EtOH, PBS, and oil, showing good agreement with the simulation results. The numerical simulations informed that the dropicle formation was influenced by the hydrophilic patches and water volume captured inside the cavity of the particle. We have observed the dimension- and time-dependent transition trend, i.e., S^1^ → T^+1d^ → R^+2d^ → C^4d^, for the dropicle formation within the 4C particles. The 4C particles captured a wide range of dropicle volumes (∼150pL up to ∼5nL), where a single 4C particle could hold up to four droplets with ∼2-7x smaller volumes compared to the state-of-the-art particles [3, 5]. Moreover, we have compared the dropicle formation within the 4C and O-shaped particles and deduced that the 4C particles could offer a much higher dropicle volume to wetting surface area ratio. Our work can inform the design of droplet volume and number per particle for developing new amplification bioassays with smaller droplets. The 4C particles with tunable dropicle configurations can lay a foundation for multiplexed high-sensitivity diagnostic assays.

## Supplementary Material

See the supplementary material for additional supporting figures with adequate descriptions.

## Acknowledgment

X.S. gratefully acknowledges the Sino-German (CSC-DAAD) Postdoc Scholarship Program, 2022 (57607866).

## Supporting information

### Numerical simulations to predict particle shape

We obtained the flow streamlines at the outlet of the microfluidic device for predicting the F_1_, F_2_, F_4_, F_9_, F_12_, and F_16_ particles’ cross-section shapes using the single-phase (**Figure S1**A) and two-phase (Figure S1B) laminar flow modules in COMSOL Multiphysics. In the numerical results, the predicted shapes of the F_1_, F_2_, and F_4_ particles had some differences, whereas the F_9_, F_12_, and F_16_ particle shapes were similar.

Apparently, the simulated particle shapes using the single-phase module were similar to the experimentally measured particles in EtOH, while the two-phase flow simulation results were closer to the particles in PBS. However, it should be noted that the predicted particle shape corresponds to the cross-sectional shape of the flow streams in the liquid state at the outlet of the microfluidic device. Once the precursor streams are cured under UV exposure, the solid particle shape will have slight differences from the predicted shape in the numerical results. Moreover, the cured particles were later washed with EtOH and transferred to PBS for experimental imaging. The media exchange also affected the particle shape. In conclusion, the two-phase flow simulation should have a more realistic prediction of the particle shape, as this model accommodates the variable viscosities of the co-flowing streams. However, the numerical and experimental results should be compared with caution and should not be expected to strictly match.

**Figure S1.**
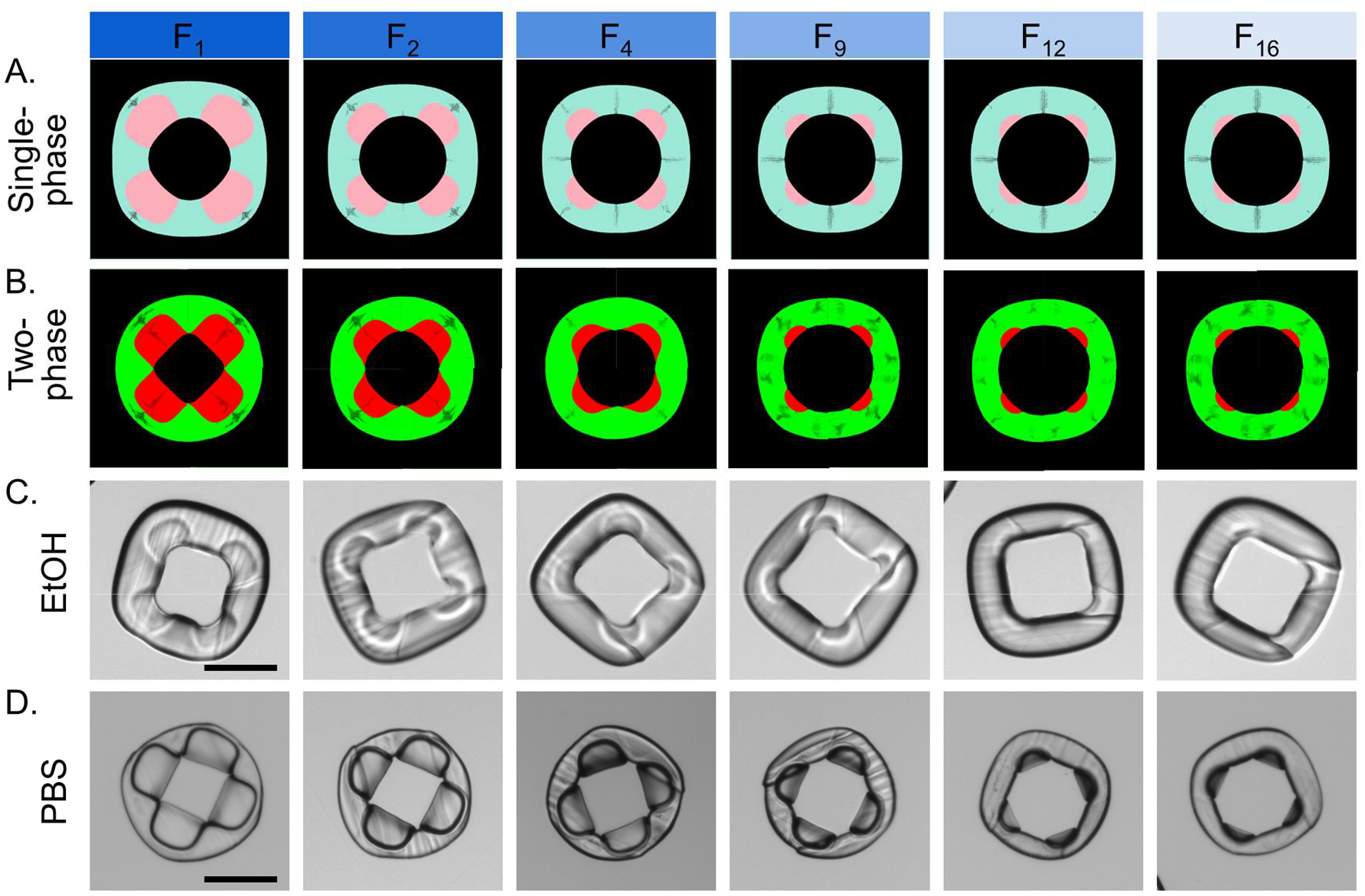
Simulated cross-section of the 4C particles using (A) single-phase and (B) two-phase flow modules. Experimentally observed particles in (C) EtOH and (D) PBS. Scale bar: 200µm.

### Numerical simulations of dropicle formation using adaptive meshing

We used “Adaptive Meshing” for modeling the dropicle formation inside a 4C amphiphilic particle (**Figure S2**). We generated a “physics-controlled mesh” with an “extremely coarse” element size (Mesh 1). We generated additional Meshes 2-11 based on Mesh 1 using “Adaptive Meshing”. Here, we set an “error estimation parameter” under Study → Solver Configurations → Solution 1 (sol1) → Time Dependent Solver → Adaptive Mesh Refinement as: “sqrt(comp1.phipfx^2 + comp1.phipfy^2 + comp1.phipfz^2)”, where comp1.phipfx, comp1.phipfy, and comp1.phypfz denoted the x, y, and z components of the phase field variable (*Θ*_*pf*_). This “error estimation parameter” refined the moving mesh along the water-oil interface. A zoomed-in Mesh 3 showed the adaptive mesh within the particle cavity. A refined mesh followed the water-oil interface with varying *T*\*. In this particular example shown in Figure S2, four individual droplets were formed at the corners of the particle and one at the center eventually, following the dropicle transition trend of S^1^ → T^+1d^ → R^+2d^ → C^4d^. One can clearly follow the mesh refinement along the partitioned droplets at the four corners and at the center of the cavity.

**Figure S2.**
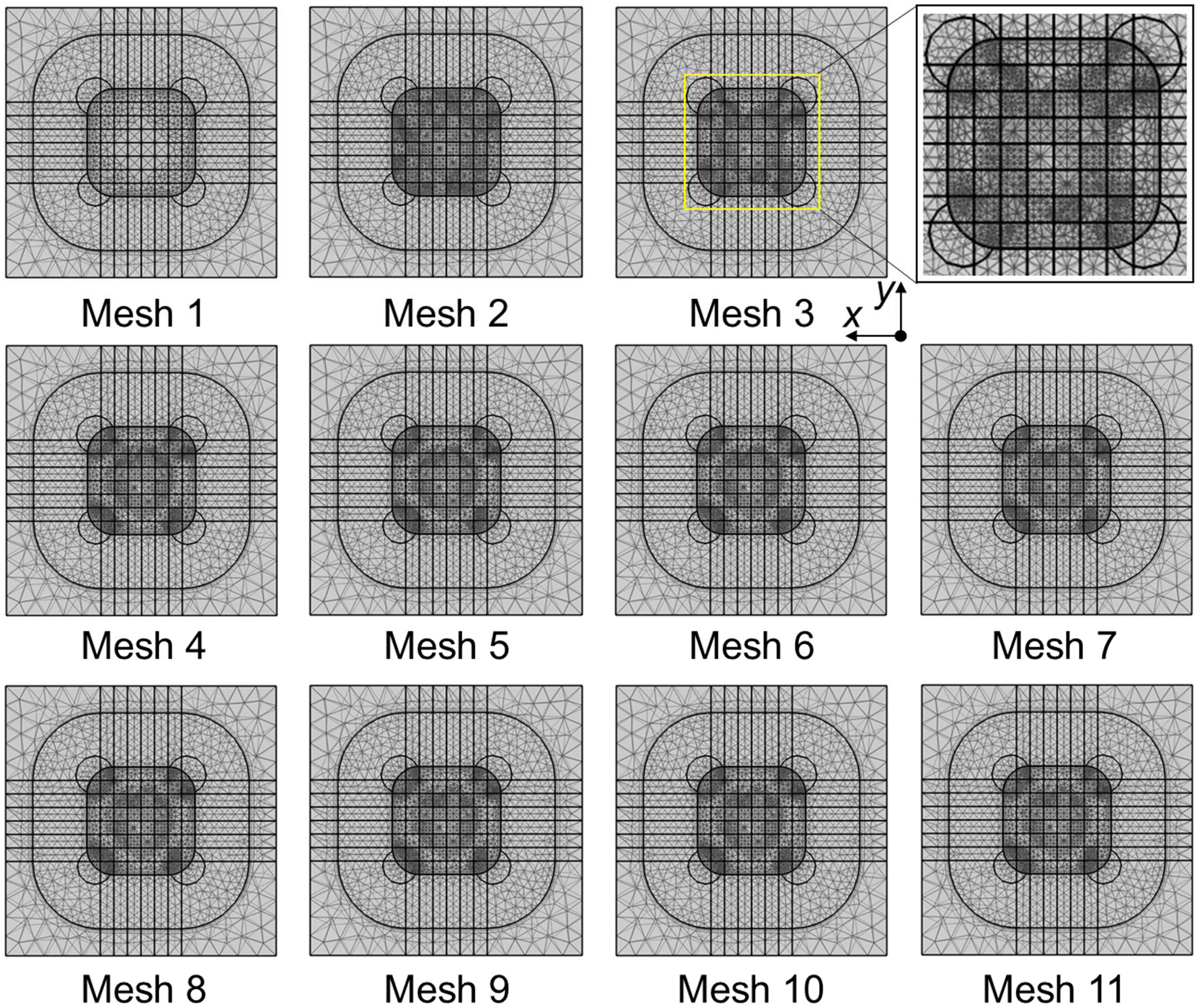
Adaptive meshes varying with *T*\* (0-1) for the dropicle formation with *H*_*d*_/*H*_*c*_ = 0.25, and *R*_1,2_, *R*_3_, *R*_4_ = 35µm, 33µm, 32µm.

### Additional experimental results on dropicle formation

**Figure S3A** depicts the F_2_ and F_4_ dropicle formation, where the particles in EtOH and PBS, and the dropicles in oil (bright field and fluorescent) can be seen. We obtained S^1^ (∼40%) and T^+1d^ (∼60%) dropicles within both particles (Figure S3B).

**Figure S3.**
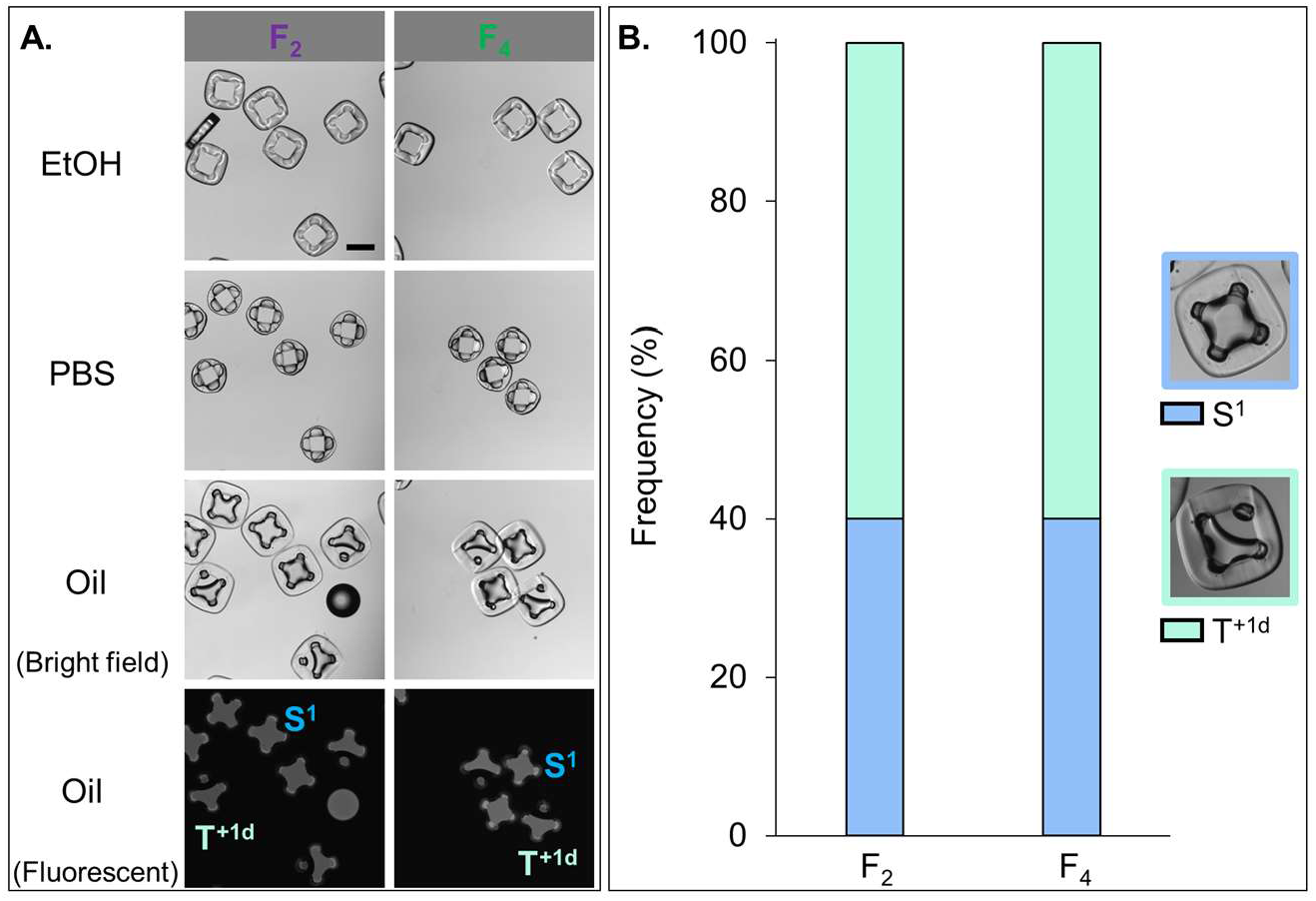
F_2_ and F_4_ dropicle formation. Scale bar: 200µm.

### Movies captions

**Movie S1**. Numerical simulations of S^1^, T^+1d^, R^+2d^ and C^4d^ dropicles formed within a 4C particle with large, medium, medium, and small size hydrophilic patches, and *H*_*d*_/*H*_*c*_ of 0.75, 0.55, 0.35, and 0.25, respectively.

**Movie S2**. Experimental dropicle formation within F_9_ particles captured for ∼45min. The dropicle configuration changed from S^1^ to T^+1d^.

**Movie S3**. Experimental dropicle formation within F_16_ particles captured for ∼3h as the dropicle configuration transitioned from S^1^ to T^+1d^ to R^+2d^ to C^4d^.

## Notes

### Competing Interest Statement

The authors have declared no competing interest.

